# Ontoclick: a web browser extension to facilitate biomedical knowledge curation

**DOI:** 10.1101/2021.03.04.433993

**Authors:** Anthony Xu, Aravind Venkateswaran, Lianguizi Zhou, Andreas Zankl

## Abstract

Knowledge curation from the biomedical literature is very valuable but can be a repetitive and laborious process. The paucity of user-friendly tools is one of the reasons for the lack of widespread adoption of good biomedical knowledge curation practices. Here we present Ontoclick, a web browser extension that streamlines the process of annotating a text span with a relevant ontology term. We hope this tool will make biocuration more accessible to a wider audience of biomedical researchers.

Ontoclick is freely available under the GPL-3.0 license on the Chrome Web Store and on the Mozilla Add-Ons for Firefox Store. Source code and documentation are available at: https://github.com/azankl/Ontoclick

## Introduction

The Human Phenotype Ontology (HPO) includes over 10,000 terms to describe the clinical features of patients with rare diseases and has proved its utility in areas that require computational processing of phenotype data such as clinical genomics (Köhler et al., 2016). However, annotating a biomedical text such as a case report in the medical literature with HPO terms remains a laborious process. Typically, a curator would identify a text span that represents a clinical phenotype (e.g. ‘abnormally large head’) and then search for this term on the HPO website. As it’s not always clear what the correct HPO term is, this often requires some trial and error. For example, searching for ‘abnormally large head’ returns no results, while searching for ‘large head’ leads to the term ‘HP:0000256 macrocephaly’. Studying the term’s definition and list of synonyms provided by the HPO website reveals that this is the correct term (macrocephaly is medical jargon for an abnormally large head). Depending on the curation workflow, the curator might then copy the term identifier (HP:0000256), the term label (‘macrocephaly’) or both into a spreadsheet or other application. The curator might also have to manually copy the associated text span from the original text (‘abnormally large head’) and a source text identifier (e.g. the Pubmed ID) into this spreadsheet or other application to complete the annotation.

## The Ontoclick Browser Extension

To facilitate this process, we have developed Ontoclick, an extension for the Chrome and Firefox web browser. Ontoclick lets the user highlight a piece of text and then tries to find a matching HPO term for it. Matching terms are returned to the user for review (Fig. 1).

**Figure 1.**
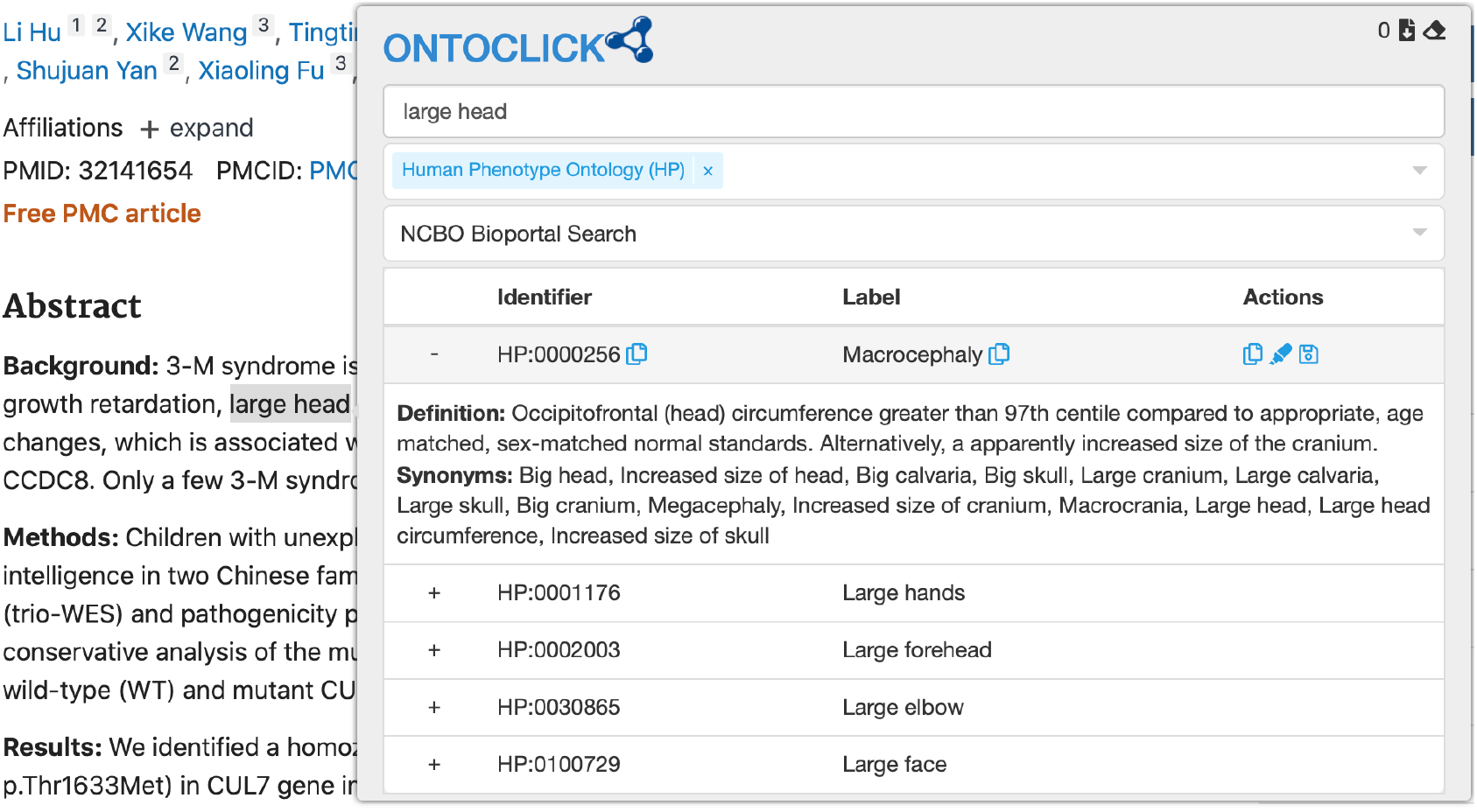
The Ontoclick popup showing HPO terms matching the highlighted text span.

Ontoclick can display a term’s definition and synonyms and provides multiple options for copying the chosen term’s identifier and/or label, with or without the highlighted text span. The copied information can be stored on the system clipboard, ready to be pasted into a spreadsheet or other application. Alternatively, multiple annotations can be stored in an internal list and exported in CSV format. Each row in the CSV file represents an annotation with separate columns for the highlighted text span, term ID and term label. The first row of the file contains the URL of the page on which the annotations were created. If the annotations were created on an abstract on the NCBI Pubmed website, the first row will contain the Pubmed ID of the annotated abstract.

In addition to the HPO, Ontoclick also allows searching for matching terms in other ontologies. We currently support searches against the Gene Ontology^1^, Mondo Disease Ontology^2^, Orphanet Rare Disease Ontology^3^ and Disease Ontology^4^.

Ontoclick uses the NCBO BioPortal Search API^5^ by default, but searches can also be performed with the NCBO BioPortal Annotator API, the EBI Ontology Lookup Service Search API^6^ and the HPO Search API^7^.(the latter two only support searches against the Human Phenotype Ontology).

Detailed installation and usage instructions are available at: https://github.com/azankl/Ontoclick

## Conclusion

We believe Ontoclick is a quick and user-friendly way to annotate biomedical texts on the web with ontology terms. While the current version is mostly geared at the HPO, the tool can be easily adapted to other ontologies. We hope that streamlined curation tools like Ontoclick will lead to more widespread adoption of biomedical curation in biomedical research.

## Footnotes

1 http://geneontology.org/

2 https://mondo.monarchinitiative.org/

3 http://www.orphadata.org/cgi-bin/index.php

4 https://disease-ontology.org/

5 http://data.bioontology.org/documentation

6 https://www.ebi.ac.uk/ols/docs/api

7 https://hpo.jax.org/app/

